# sandwrm: an R package for estimating Wright’s neighborhood size and species-level genetic diversity

**DOI:** 10.1101/2025.05.19.654925

**Authors:** Zachary B. Hancock, Gideon S. Bradburd

**Affiliations:** Department of Ecology & Evolutionary Biology, University of Michigan, Ann Arbor, MI 48103

**Keywords:** isolation by distance, continuous space, effective population size, Bayesian

## Abstract

In most natural populations, individuals in close proximity are more related on average than those at greater distances; this pattern gives rise to geographic population genetic structure. Despite extensive theoretical work on spatial population genetics, few empirical methods exist to estimate the components of theoretical models of genetic relatedness in continuous space. One classic model of relatedness in continuous space is the Wright-Malécot model, which predicts that the probability of identity-by-descent decays as a function of geographic distances. The shape of this decay curve is dictated by the dynamics of local dispersal and mating, as well as population density. This model can be reformulated to describe the probability of identity-by-state, in which case it decays to an asymptote, the value of which is determined by the historical demography of the population. Collectively, these features can be modeled in a likelihood-based framework to estimate neighborhood size and long-term diversity from pairwise genetic and geographic distance. In this article, we introduce the R package `sandwrm` Spatial Analysis of Neighborhood size and Diversity using WRight-Malécot), which takes a Bayesian approach to estimate key parameters of populations that are both dispersal-limited and distributed continuously across a landscape.

## 1 Introduction

Many organisms in nature are dispersal-limited and tend to be more related to individuals nearby than to those far away. This pattern, known as isolation-by-distance (IBD) (Wright, 1943), has been widely documented in a diversity of organisms (e.g., Aguillon et al., 2017; Ramachandran et al., 2005; Rousset, 1997; Sharbel et al., 2000), and reflects the reality that most organisms move, forage, compete, and locate mates in continuous geographic space (Battey et al., 2020; Bradburd and Ralph, 2019). However, most population genetic inference programs that have been used in empirical studies treat organisms as well-mixed, discrete populations connected by occasional migration (e.g. Alexander et al., 2009; Beerli et al., 2019; Epperson, 2003; Pritchard et al., 2000). Past work has shown that many population genetic summary statistics, like Watterson’s *θ* and Tajima’s *D*, as well as estimates of population genetic structure, are biased in the presence of continuous structure (Battey et al., 2020; Bradburd et al., 2018; Frantz et al., 2009). In addition, continuous population structure can lead genome-wide association studies (GWAS) to identify spurious genetic associations with completely environmentally-determined traits (Battey et al., 2020).

While most popular inference programs utilize models that are non-spatial, theoretical population genetics has a long history of describing populations in continuous space. The first of these models was introduced by Sewall Wright (1943; Wright 1946) and Gustave Malécot (1948); they imagined a homogeneous, infinite landscape in which individuals disperse in two dimensions following a bivariate Gaussian distribution with a mean of zero and standard deviation in each dimension of *σ*, which is interpreted as the “dispersal distance.” Elaborating on the Wright-Malécot model, Barton et al. (2002) formalized the probability that individuals are identical-by-descent, *F*_*i,j*_, given *σ* and the geographic distance, *d*_*i,j*_, separating individuals *i* and *j* as:

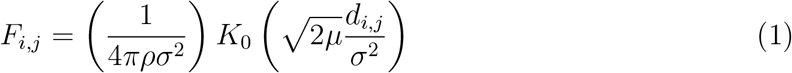

where *ρ* is the population density, *π* is the mathematical constant, *µ* is the mutation rate, and *K*_0_ is a modified Bessel function of the second kind (Barton et al., 2002; Malécot, 1948; Ringbauer et al., 2017; Wright, 1943). A key value in this expression is the compound parameter 4*πρσ*^2^, which is known as “Wright’s neighborhood size” (𝒩). This parameter captures the number of potential breeding individuals within a radius defined by the per-generation parent-offspring dispersal distance, *σ*. Thus, 𝒩 captures the local rate of genetic drift. Using simulations, Hancock et al. (2024) demonstrated that estimates of 𝒩 are relatively insensitive to demographic shifts that occurred long ago, and tend to reflect contemporary population dynamics. Furthermore, for two-dimensional systems, it determines the slope of the isolation-by-distance curve, and thus informs us about the scale of spatial autocorrelation of genetic ancestry.

While 𝒩 captures local demographic and evolutionary dynamics that might have occurred in the recent past, researchers are often also interested in long-term, global trends across an entire metapopulation, or even a species’ range. These trends can inform us about ancient population bottlenecks or expansions, inbreeding, and a species’ adaptive potential (Charlesworth, 2009; Mable, 2019; Willi et al., 2006). While the slope of the isolation-by-distance curve in two-dimensional habitats is shaped by 𝒩, the long-term inbreeding effective population size (*N*_*e*_) is reflected by the asymptote of pairwise sequence dissimilarity against geographic distance. An estimator of this long-term value was introduced by Hancock et al. (2024) and termed “collecting-phase *π*” (*π*_*c*_), so named because it captures the outcome of demographic processes from the collecting phase of the coalescent (Wilkins, 2004). This parameter can be thought of as the diversity of a spatially over-dispersed sample of individuals (i.e., not biased by spatial autocorrelation of ancestry). See Hancock et al. (2024) for additional details.

Historically, 𝒩 has been estimated almost exclusively by regressing linearized F*ST* against geographic distance, a method introduced by Rousset (1997). Recently, Hancock et al. (2024) introduced a Bayesian statistical approach for jointly estimating neighborhood size (𝒩) and long-term diversity (*π*_*c*_) from pairwise genetic and geographic distance data. In this paper, we present `sandwrm` (Spatial Analysis of Neighborhood size and Diversity using WRight-Malécot), a user-friendly R-package for the analysis of spatial genetic data. Specifically, `sandwrm` employs the model from Hancock et al. (2024) to estimate Wright’s neighborhood size and *π* in the collecting phase, and generates diagnostic plots for ease of visualizing parameter estimates and evaluating model fit.

## 2 Usage

The `sandwrm` package can be installed from the github repository https://github.com/zachbhancock/sandwrm, where users will also find installation instructions and a basic tutorial. Below, we briefly describe how `sandwrm` can be used, including a discussion of data preparation, analysis choices, and explanation of model output and interpretation.

The `sandwrm` package is coded in the R programming language (R Core Team, 2022) wrapped around an inference engine implemented in `rstan` (Stan Development Team, 2025). Details of the model are described in Hancock et al. (2024), but will be expounded upon briefly here.

### 2.0.1 Model

We model the pairwise sample homozygosity 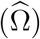, which we define as the the complement of pairwise diversity, i.e., 1-*D*_*XY*_, as a draw from a Wishart distribution:

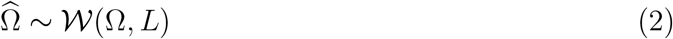

where *L* is the degrees of freedom of the Wishart (which we discuss further below), and Ω is a parametric covariance that is a function of geographic distance and the parameters of our model. Specifically,

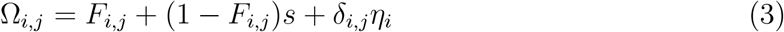

where *s* = 1 − *π*_*c*_ (that is, 1 minus *π* in the collecting phase), *δ*_*i,j*_ is the Kronecker *δ*, and *ηi* describes inbreeding specific to the *i*th individual (Hancock et al., 2024; Ringbauer et al., 2017). The quantity *F*_*i,j*_ is the probability of identity by descent between samples *i* and *j*, and is defined as a function of the geographic distance, *d*_*i,j*_, between them:

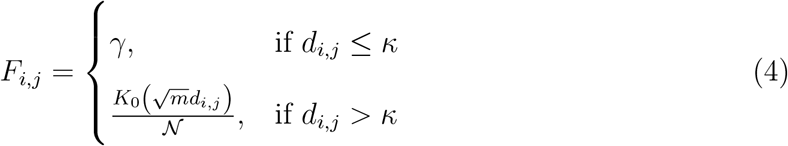

where *m* is a compound parameter equal to 2*µ/σ*^2^, *γ* is the rate of coalescence between nearby individuals, and “nearby” (i.e., within distance *κ* of each other) is defined as near enough that they can be considered to be randomly mating (see Hancock et al. (2024) for an additional discussion).

### 2.0.2 Data preparation

To execute the model, users specify a pairwise genetic distance matrix. The genetic distance calculated should be pairwise *π*, also known as *D*_*XY*_ – the average pairwise sequence dissimilarity per base pair. Note that the appropriate denominator in calculating *π* is the total number of basepairs genotyped in both samples being compared. Pairwise *π* calculated solely from polymorphic sites should not be used in this analysis. An avenue for researchers to acquire a pairwise *π* matrix is to generate it using `pixy` (Korunes and Samuk, 2021), which accepts a VCF file that includes polymorphic and monomorphic sites. Researchers who assembled restriction-associated DNA (RAD) tags or other short-read datasets can generate a VCF file with included monomorphic sites using `STACKS` (Rochette et al., 2019) with the call `–vcf-all` in the `populations` function. From this, `pixy` will estimate pairwise *π*, which can be converted to a matrix for the `sandwrm` analysis. Additional documentation on how to prepare a VCF file that contains monomorphic and polymorphic sites can be found in the `pixy` documentation. For researchers with low-coverage, whole genome data, pairwise genetic distance matrices can be acquired from methods such as `distANGSD` (Zhao et al., 2022).

Users must also specify a pairwise geographic distance matrix. Pairwise geographic distance can be in whatever units the user wishes (though note that estimated neighborhood size is in units of individuals per area, so the units chosen will affect the estimated value of 𝒩). For sampling over large geographic regions, users may wish to calculate great-circle distance; for sampling on smaller spatial scales, users may wish to use distances based on, e.g., Universal Transverse Mercator (UTM) coordinates, which are more accurate over smaller scales.

In addition to pairwise genetic and geographic distances, there are two other quantities that users must specify in order to initiate a `sandwrm` analysis: the number of independent loci in their dataset (*L*) and the spatial cutoff within which the model of isolation by distance no longer applies (*κ*).

The parameter *L* determines the degrees of freedom of the Wishart distribution used to model the data; larger values indicate higher confidence and concomitantly lower acceptable “noise” in inference. In principle, *L* is the number of independent (i.e., unlinked) loci used to estimate pairwise genetic relatedness. In practice, it is often unclear precisely what value to specify for *L* because loci may be in linkage disequilibrium (LD) over long stretches of the genome. Thinning SNPs for LD (e.g., by retaining only a single SNP per RAD locus) may not completely remove SNPs that are evolving nonindependently, which can result in an inflated confidence due to pseudoreplication. In addition, *L* only captures the uncertainty associated with estimates of pairwise homozygosity; it does not capture uncertainty in the Wright-Malécot model itself (i.e., uncertainty due to stochasticity in the genealogical and geographic processes that generate the observed distribution of individuals and their relatedness). As a result, very large values of *L* made possible by, e.g., whole-genome resequencing datasets, could lead to unrealistically low uncertainty. We recommend that researchers test a range of values of *L* below that obtained after thinning for LD.

The units of *κ* are the same as the distance matrix (e.g., if the geographic distance is measured in kilometers, the dispersal distance is in units of kilometers). Importantly, if the random-mating distance is unknown, this value can be set arbitrarily low without significantly affecting the model performance, but if it is set too high it might deplete the signal of isolation-by-distance, especially if the signal of correlation between ancestry and geographic distance decays over very short distances.

To begin a `sandwrm` analysis, users specify these quantities (genetic distance, geographic distance, *L*, and *κ*) as arguments to the `prepareData` function, which will generate an R object that can then be passed to the function `sandwrm`.

The function `sandwrm` initiates a Markov chain Monte Carlo (MCMC) analysis that characterizes the posterior distribution of all model parameters, including 𝒩 and *π*_*c*_. Users will need to specify how many independent MCMC chains they would like to run and how many steps for these chains to take. We recommend at least two independent chains to confirm that both are converging on the same posterior distribution. The Hamiltonian Monte Carlo algorithm implemented in `rstan` is very efficient, so most runs of this model are able to mix effectively in 5,000 iterations or fewer. A typical run would look something like:

~~~
sandwrm(dataBlock=dataBlock,
        nChains=4,
        nIter=2000,
        prefix=“test”)
~~~

By default, a successful call to `sandwrm` analysis generates two output files: an R object saved in binary file format, and a PDF that includes various figures, both of which will have a file name determined by the “prefix” argument specified. The first of these figures displays trace plots of the MCMC to visually inspect for convergence (Figure 1). The second is a series of panels showing the joint marginal distributions of all pairs of estimated parameters (Figure 2). The last plot shows the fit of the model to the empirical data (Figure 3).

**Figure 1:**
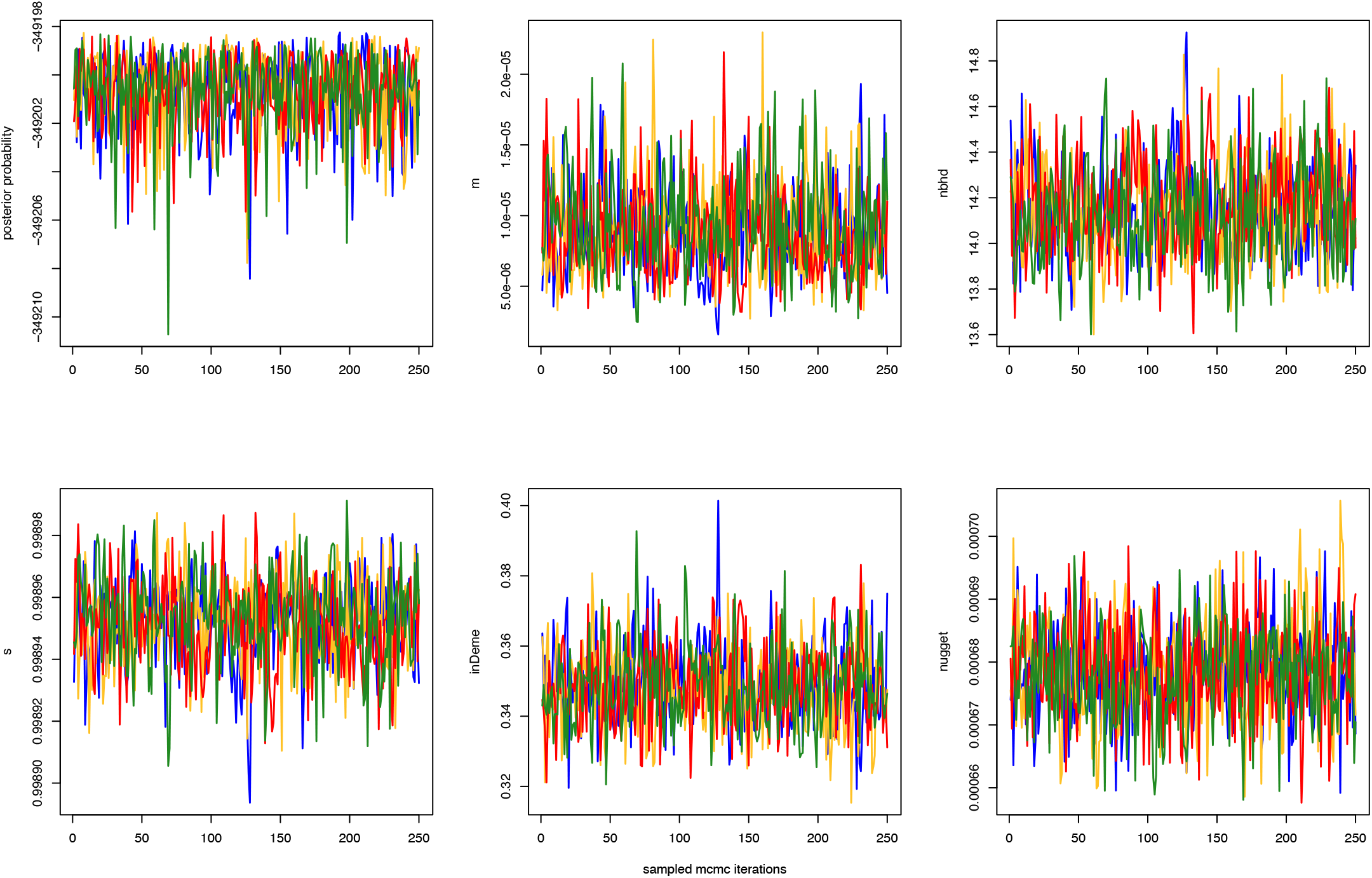
MCMC trace plots produced by `sandwrm` for the posterior probability and each estimated parameter. Different colors represent the four independent MCMC chains.

**Figure 2:**
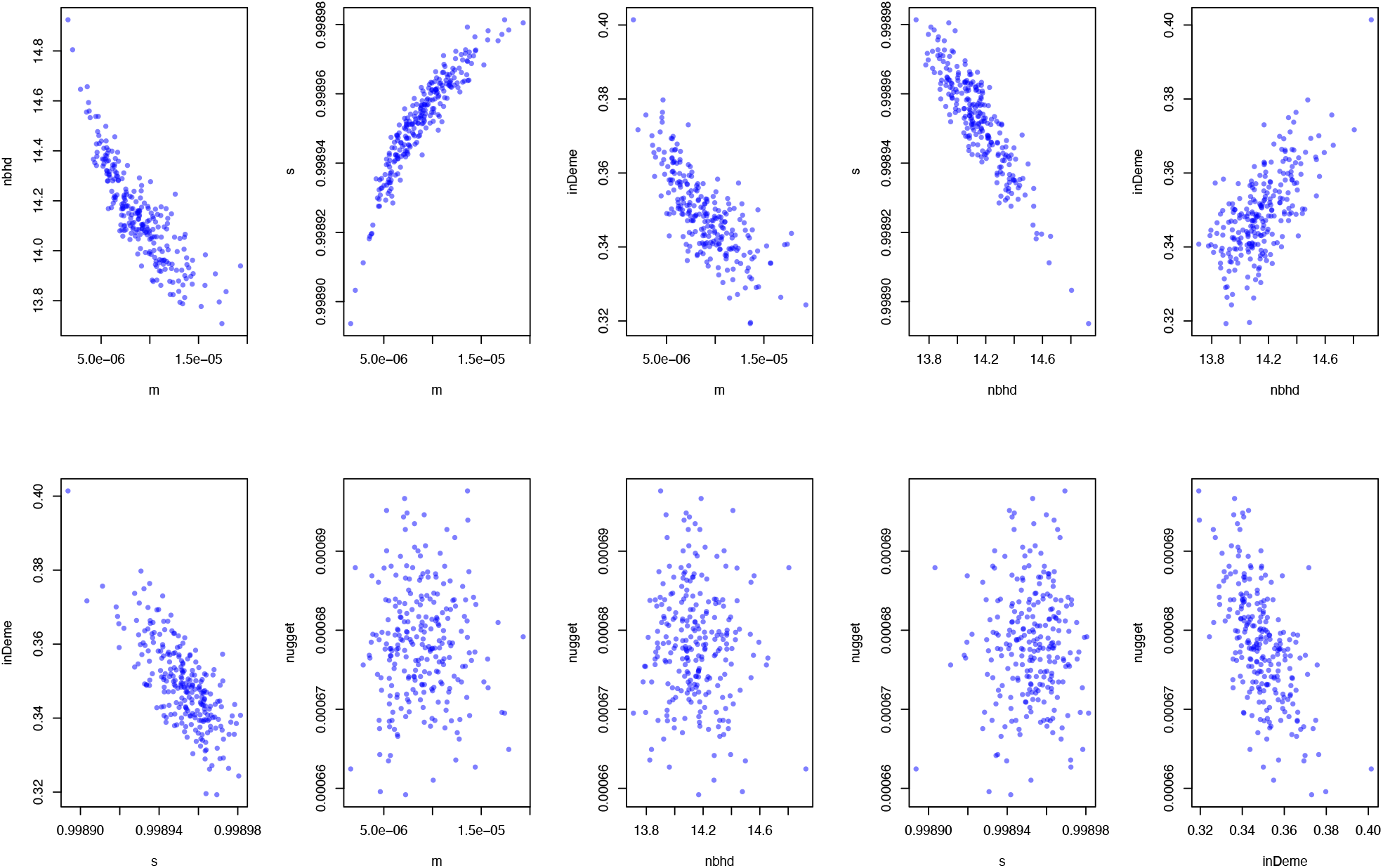
Example output of the correlations panels between estimated parameters in `sandwrm`. The function will produce figures for each independent MCMC chain with the colors matching those from Fig. 1.

**Figure 3:**
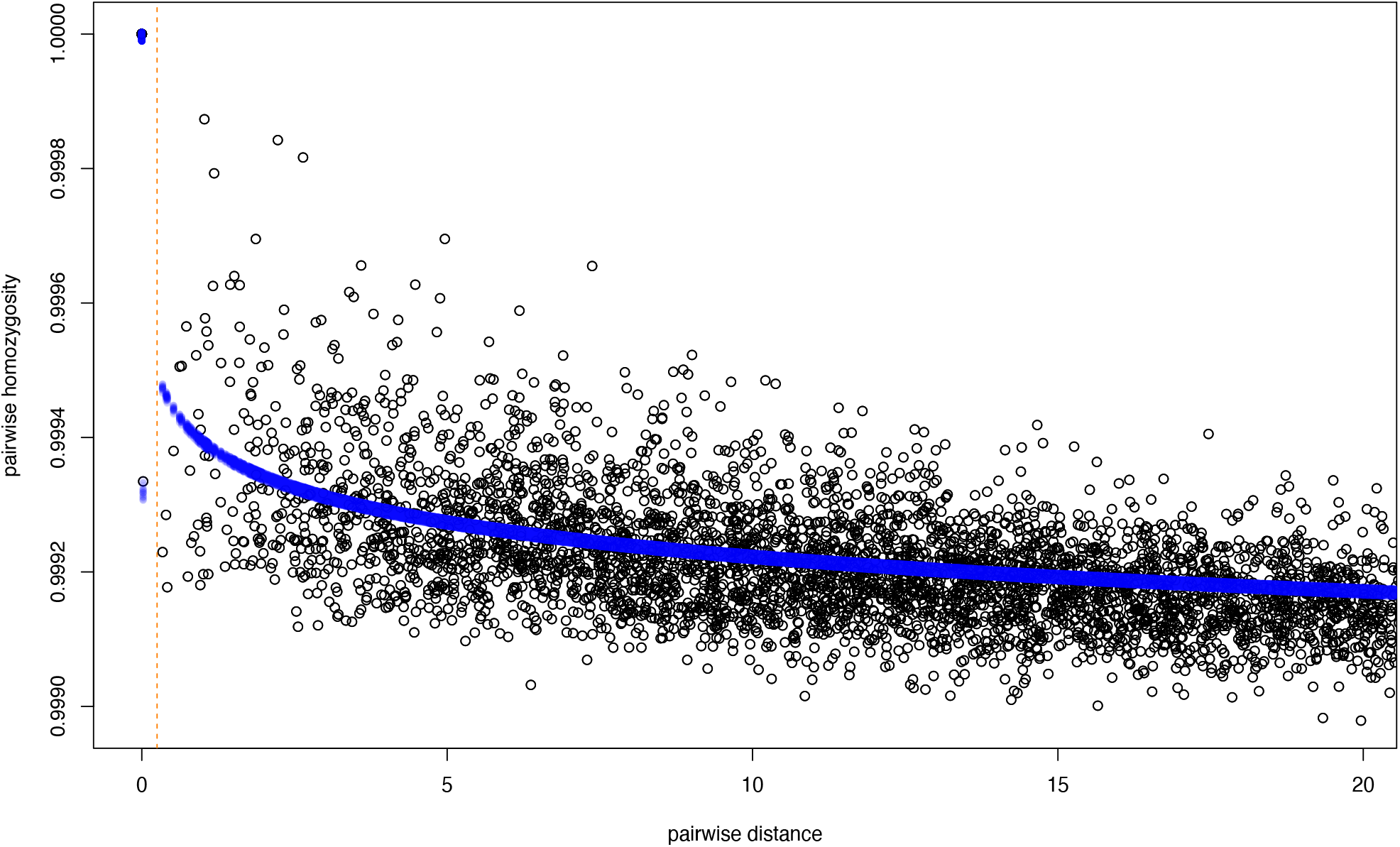
Model fit output from a single chain of a `sandwrm` run. Dark circles are the empirical estimates, blue circles are the fit of the model. The dashed line is *κ*.

The output R object file includes all the estimated parameters from each independent MCMC chain. Each of the parameters and their abbreviations are listed in Table 1.

**Table 1.**
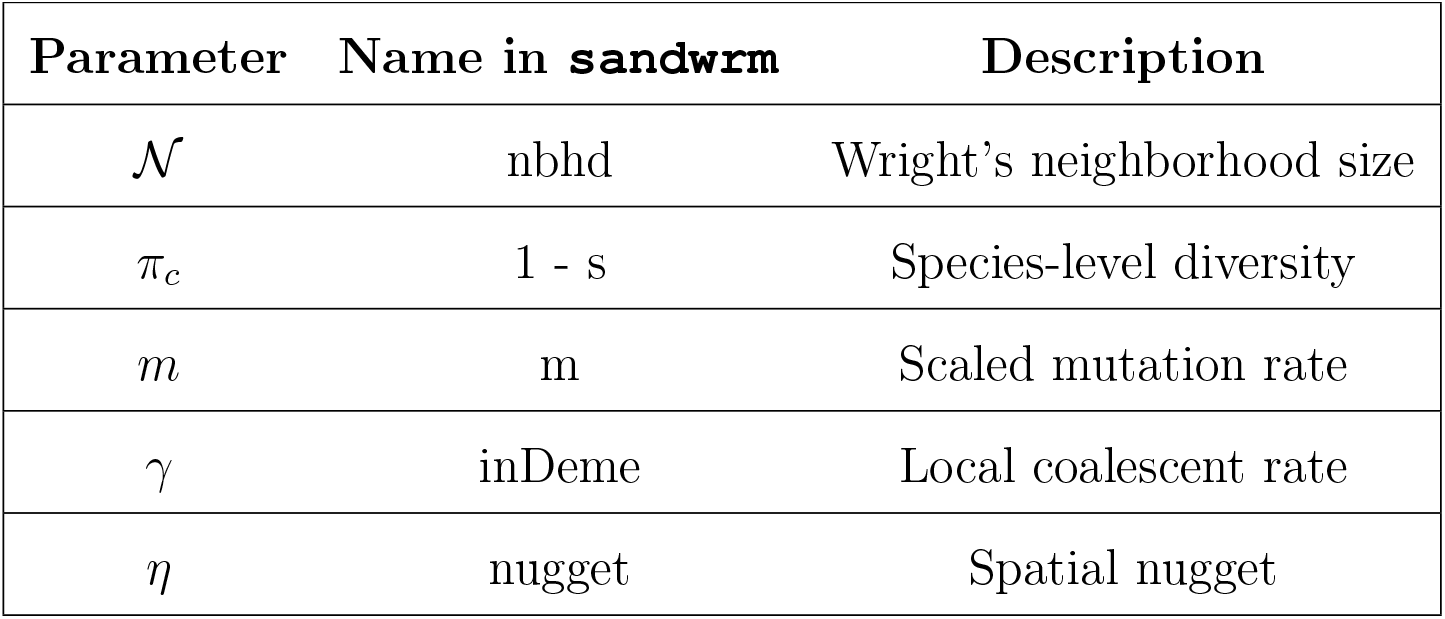

To establish benchmarks for model runtime, we recorded the computational run-time for a single MCMC chain for datasets consisting of 25, 50, or 100 individuals (*N*) and of 1,000, 10,000, or 100,000 independent loci (*L*). The test dataset used for benchmarking is the same as that used in the “Example workflow” section below. To calculate runtime, we used the package `tictoc` (Izrailev, 2025). The benchmarking results are displayed in Figure 5. Across the tested datasets, *L* had a minimal impact on run-time, whereas increasing sample size led to an approximately cubic increase in run-time. This faster-than-linear increase in run-time with sample size is unavoidable, as the likelihood calculation for the Wishart distribution involves inverting a covariance matrix, an operation that has a time complexity of *O*(*n*^3^).

## 3 Example workflow

In this section, we provide an example workflow. The dataset for this example is included within the `sandwrm` package as

~~~
example_locs.txt example_pwp.txt
~~~

This is a simulated dataset of a continuous-space model in the forward-time simulator `SLiM` v.3.6 (Haller and Messer, 2019) as described in Hancock et al. (2024). The locations file contains coordinates for each individual and can be converted into a distance matrix as follows:

~~~
   data(“example_locs”, “example_pwp”)
   library(fields)
   coords <- example_locs[,c(“x”,”y”)]
   geoDist <- rdist(coords)
~~~

Next, the function `prepareData` can be used to construct the data block, which is a list of several components, that is required to run `sandwrm` The function is used as follows:

~~~
   L <- 1e4 #number of independent loci
   k <- 0.25 #kappa parameter
   dataBlock <- prepareData(genDist = example_pwp, geoDist = geoDist, L = L, k = k)
~~~

With the data block constructed, the `sandwrm` is called as indicated above. From the out.Robj file, we can extract and generate a histogram of estimated 𝒩 using the `plyr` and `dplyr` packages as follows:

~~~
   load(“out.Robj”)
   nbhd <- rstan::extract(out$fit, “nbhd”,
   inc_warmup=TRUE, permute=FALSE)
   #make long
   nbhd_long <- plyr:adply(nbhd, c(1, 2, 3)) %>%
   dplyr::rename(“iteration”=“iterations”,
   “chain”=“chains”,
   “nbhd”=“V1”) %>%
   mutate(chain = gsub(“chain:”, ““, chain)) %>%
   dplyr::select(-parameters)
   #plot nbhd
   hist(nbhd_long$nbhd)
~~~

The remaining parameters can also be extracted and plotted in this way (Figure 4).

**Figure 4:**
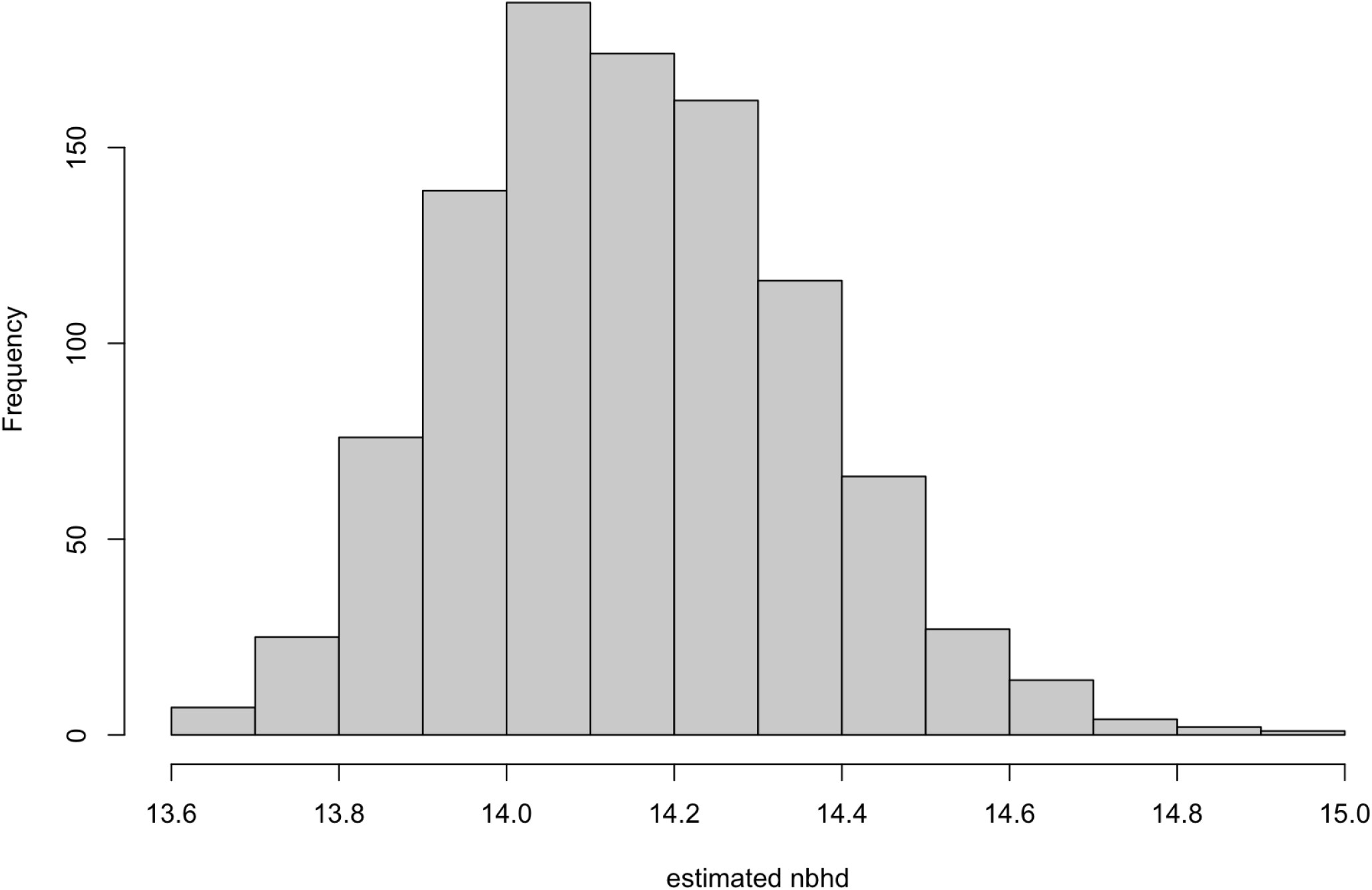
Histogram of 𝒩 estimated from the `sandwrm` example dataset across all four chains.

**Figure 5:**
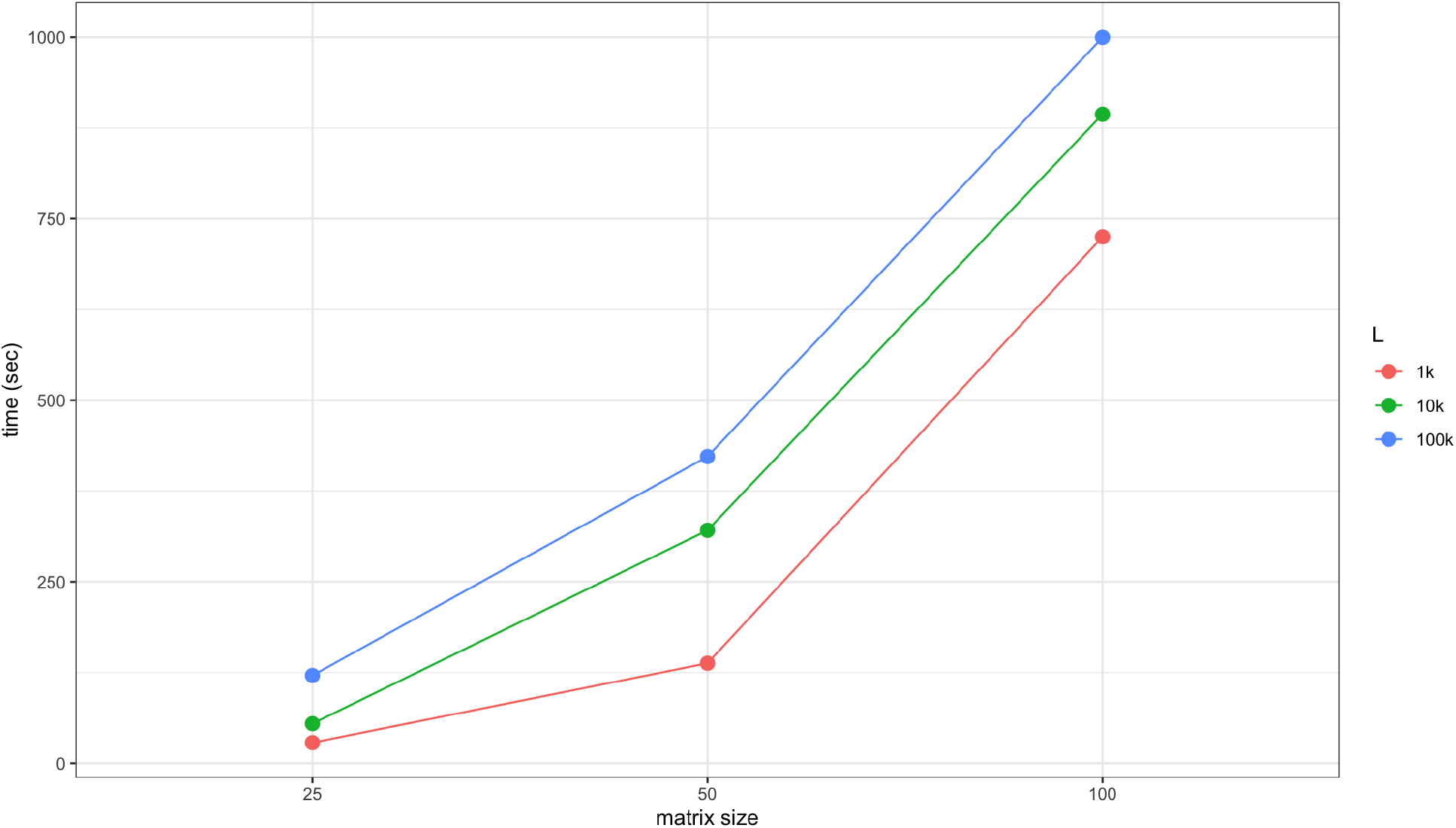
Benchmarking results for matrix sizes of 25, 50, and 100 across three different values of *L*: 1000, 10000, and 100000.

## 4 Conclusion

Despite the ubiquity of organisms in nature that are both dispersal-limited and continuously distributed across the landscape, relatively few methods exist for using DNA sequence data to learn about important theoretical parameters like Wright’s neighborhood size. For methods that do exist, even fewer have been packaged in such a way as to permit general use among empiricists. In this article, we describe a new R package, `sandwrm`, intended to remedy this issue by providing users with an easy, straightforward tool for estimating both 𝒩 and a novel statistic that captures genetic diversity at the species-level and is unbiased by local structure, *π*_*c*_. We have provided a general workflow for how to use the package; additional tutorials can be found at https://github.com/zachbhancock/sandwrm.

## 5 Acknowledgments

We’d like to thank members of the Bradburd lab, Kristen Wacker, and Seth Temple for helpful feedback on this manuscript. Research reported in this publication was supported by the National Institute of General Medical Sciences of the National Institutes of Health under Award Number R35GM137919 (awarded to GSB). The content is solely the responsibility of the authors and does not necessarily represent the official views of the NIH.

## 6 Data accessibility

The package `sandwrm` can be installed in its most up-to-date form from the github repository: https://github.com/zachbhancock/sandwrm.

